# Hi-C Analyses with GENOVA: a case study with cohesin variants

**DOI:** 10.1101/2021.01.22.427620

**Authors:** Robin H. van der Weide, Teun van den Brand, Judith H.I. Haarhuis, Hans Teunissen, Benjamin D. Rowland, Elzo de Wit

## Abstract

Conformation capture-approaches like Hi-C can elucidate chromosome structure at a genome-wide scale. Hi-C datasets are large and require specialised software. Here, we present GENOVA: a user-friendly software package to analyse and visualise conformation capture data. GENOVA is an R-package that includes the most common Hi-C analyses, such as compartment and insulation score analysis. It can create annotated heatmaps to visualise the contact frequency at a specific locus and aggregate Hi-C signal over user-specified genomic regions such as ChIP-seq data. Finally, our package supports output from the major mapping-pipelines. We demonstrate the capabilities of GENOVA by analysing Hi-C data from HAP1 cell lines in which the cohesin-subunits SA1 and SA2 were knocked out. We find that ΔSA1 cells gain intra-TAD interactions and increase compartmentalisation. ΔSA2 cells have longer loops and a less compartmentalised genome. These results suggest that cohesin^SA1^ forms longer loops, while cohesin^SA2^ plays a role in forming and maintaining intra-TAD interactions. Our data supports the model that the genome is provided structure in 3D by the counter-balancing of loop formation on one hand, and compartmentalization on the other hand. By differentially controlling loops, cohesin^SA1^ and cohesin^SA2^ therefore also affect nuclear compartmentalization. We show that GENOVA is an easy to use R-package, that allows researchers to explore Hi-C data in great detail.

## Introduction

The organization of the genome inside the nucleus can be measured using proximity ligation assays such as Hi-C(1). A detailed picture is emerging of a hierarchically organized genome. Chromosomes are subdivided into compartments or compartmental domains which form microenvironments that segregate active and inactive chromatin(2). Compartments can be further segmented into Topologically Associated Domains (TADs)(3, 4), which are genomic regions that show increased self-interaction. Although the initial observation of TADs was largely phenomenological(5), the mechanisms shaping TADs are starting to be understood. TADs are a collection of loops formed by the ring-shaped cohesin complex(6, 7).

The mechanism by which cohesin forms these loops, and by extension TADs, is loop extrusion (8). In this model, cohesin processivily increases the size of chromatin loops. Extrusion is halted when cohesin encounters the CCCTC-binding factor (CTCF) bound to DNA. The orientation of the CTCF consensus-motifs is important for the ability of CTCF to act as a boundary-element for chromatin loops (9). The majority of stable loops observed in Hi-C maps brings together CTCF motifs in opposite orientation (the ‘convergency rule’) (9, 10). We and others have shown that stabilising chromatin-bound cohesin, by depleting the cohesin-release factor WAPL, leads to more and longer loops(7, 11). These loops follow the convergency rule less strictly, and are generally extensions of wild-type loops, suggesting that loop-anchors collide in de absence of WAPL(7, 12). These observations show that by regulating the cohesin complex we can critically influence the organization of the genome inside the nucleus. The cohesin complex is a multimeric complex consisting of the core proteins SMC1, SMC3, RAD21/SCC1 and a STAG/SA subunit. There are two different cohesin variants, that contain either SA1 or its homologue SA2. Recent studies suggested that cohesin^SA1^ forms long CTCF-anchored loops(13–15), whereas cohesin^SA2^ is involved in the formation of promoter-enhancer loops(13, 16).

Many recent discoveries concerning the organisation of the 3D genome and the role of cohesin in this has been learned from Hi-C, which is an all-versus-all chromosome conformation capture method(1). Visualising individual chromatin loops requires Hi-C maps with resolutions of at least 20kb (17). Since Hi-C data is a pairwise analysis method, increasing the resolution requires a quadratic increase in reads. For this reason, Hi-C datasets are often very large. More recently, higher-resolution methods like micro-C (18) have emerged, resulting in even larger datasets. These large amounts of data call for purpose-built and highly powerful computational methods.

Several software-packages for Hi-C analysis and visualisation have been described in recent years (19). Some of these focus on generating tracks or snapshots of regions of interest (20, 21). Another powerful feature is aggregating Hi-C data on specific features like loops, also referred to as pile-ups (7, 22–25). By averaging the limited signal of many features, one can surmise general changes in nuclear organization from changes in signal distribution. These aggregations are conceptually similar to metaplots in ChIP-seq and ATAC-seq analyses. The Hi-C analysis methods referenced above are currently scattered over many packages and programming languages. This dispersed landscape of tools is cumbersome for many experimentalists, as it forces them to spend time learning how to use each of these tools and to become versed in multiple programming languages. Here we present GENome Organisation Visual Analytics (GENOVA): an R-software package for Hi-C data-analysis. This package is designed to be the one-stop shop for 3D genomics. It features all of the key Hi-C analyses and works with all major mapping-pipelines. GENOVA can be downloaded and installed from github.com/dewitlab/GENOVA.

GENOVA has previously been used to study the role of the ChAHP in nuclear organization(26), to investigate the loss of all CTCF anchored loops in a CTCF point mutant(27) and other studies(28, 29). In the current study we present GENOVA in detail and use it to chart the roles of SA1 and SA2 in genome organisation. We generated knockouts of each homolog in human HAP1 cells. GENOVA enabled the integration of published Hi-C data of knockdowns and acute depletions (13–15, 30). Using GENOVA we were able to determine the contribution of cohesin^SA1^ and cohesin^SA2^ to genome organization.

## Methods

The basic principle in Hi-C data analysis is identifying ligations between non-contiguous restriction fragments. This is achieved by performing paired-end sequencing of a Hi-C template. Hi-C mapping pipelines have the following steps in common. First, paired-end sequence reads are mapped to a reference genome. When the paired ends fall on different restriction fragments this amplicon is identified as a valid interaction pair. Next, the valid pairs are summed over equally-sized (e.g. 10 kilobase) interaction bins. Finally, the resulting contact matrix is normalized to account for biases using iterative correction (31) or matrix balancing (32). The most common pipelines (Hi-Cpro, juicer and cooler) perform these steps but produce different output formats (23, 33, 34).

### Loading and representation of Hi-C data

In GENOVA, the contact matrices are loaded into contacts-objects, which stores the matrices in a compressed sparse triplet format and the user-added metadata (e.g. colours and sample-names) of one Hi-C dataset (fig. 1A). There is also the option to calculate Z-score normalised values. These scores express data in units of standard deviation relative to other values at equal distance. This can be of use when exploring small (i.e. 1 by 1 bin) far-*cis* features, as the increase in sparsity at these distances means that it is more difficult to separate noise from true local contact-enrichment. Data from the Juicer, Cooler and HiC-pro pipelines can all be loaded with the same function inside GENOVA. The Juicer pipeline produces .hic-files that are parsed with the strawr-package. The Cooler pipeline produces “.cooler”-files that stored in the HDF5 standard. The Rhdf5-package enables the loading of these into R.

**Figure 1:**
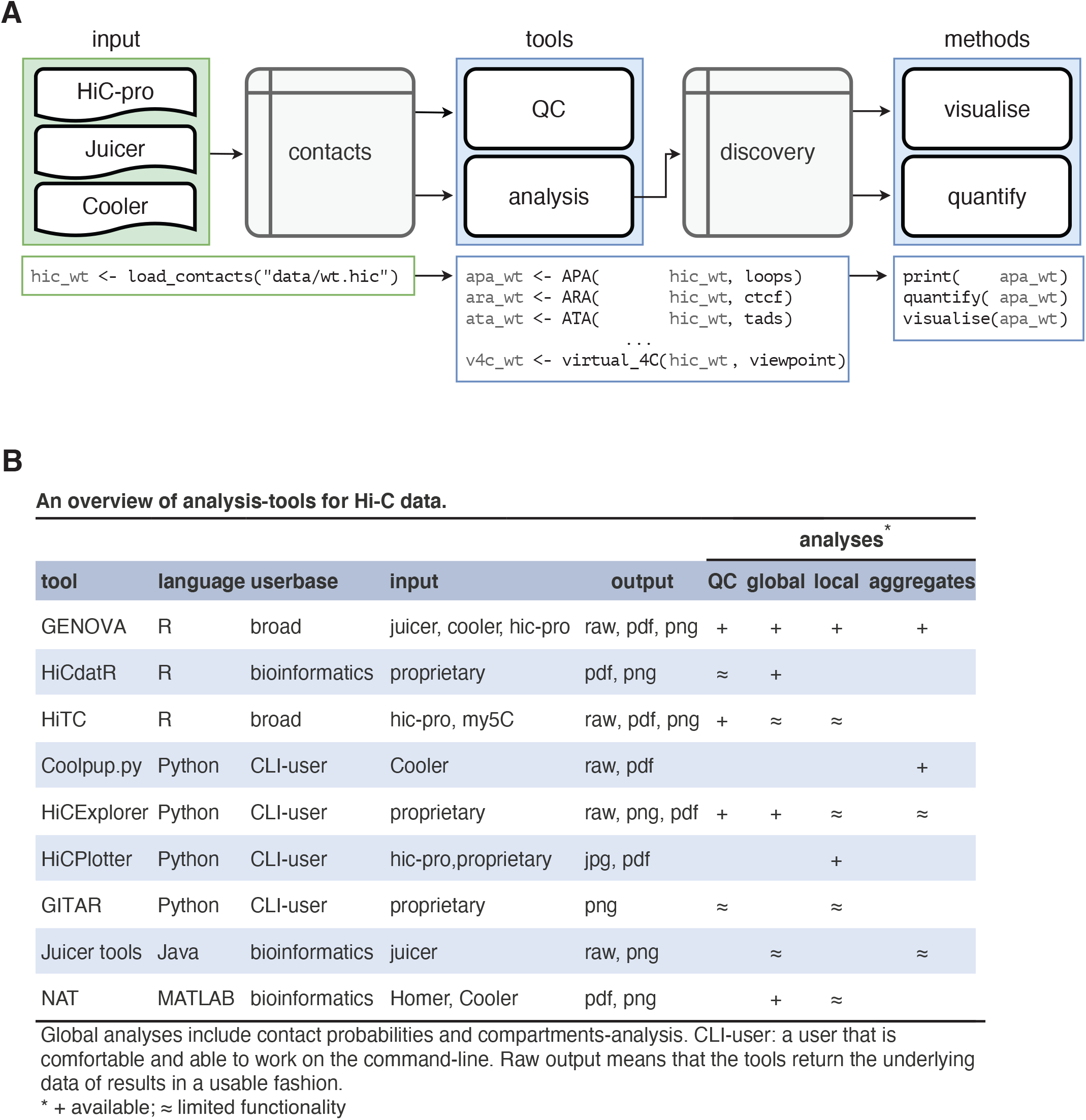
GENOVA is a pipeline-agnostic R-package and includes the majority of Hi-C analyses. A) Data from the three major pipelines can be loaded with the `load_contacts` tool into a contacts-object. Quality control and other analyses can then be performed on these objects: all tools generate the results in the form of a discovery-object. The user can print, visualise and quantify these objects. B) An overview of the tools and options in GENOVA and other Hi-C software. The majority of the available software focus on a subset of the possible analyses and are often restricted to specific mapping pipelines.

After contact-objects are made, the user can analyse these with the tools (R-functions to analyse Hi-C data) in GENOVA. All tools have a similar syntax and standardised output: the discovery object. An added benefit of using contacts- and discovery-objects is that they are portable: they contain all the information of a Hi-C dataset or result, including metadata. This averts common errors, like swapping labels, and facilitates sharing (raw) data of analyses with collaborators. The user can visualise the discovery-objects, as well as quantify them for further analysis (fig. 1A).

The main benefit of using GENOVA is that it comprises a large set of available tools, that are otherwise distributed over a number of different software packages and programming languages. The tools in GENOVA can perform quality-control, generate tracks, visualize contact matrices and aggregate Hi-C data over genomic features (fig. 1B). This has resulted in a package that can be used to run the majority of analyses currently used in the literature within a single programming environment. We will discuss these tools in detail below.

### Quality control

The first analysis-step after loading the data is to perform quality control to check the integrity of the Hi-C experiment. A good indicator of the quality of a Hi-C library is the percentage of reads mapping in *cis*. Previous work has shown that the expected amount of intra-chromosomal contacts is in the 90-93% range in both mouse embryonic stem cells and in human K562 cancer cells (35). This implies that the desired *cis*-percentage should also be in this range. To test this, users can run the `cis_trans`-tool, which computes this percentage genome-wide (fig. 2A).

**Figure 2:**
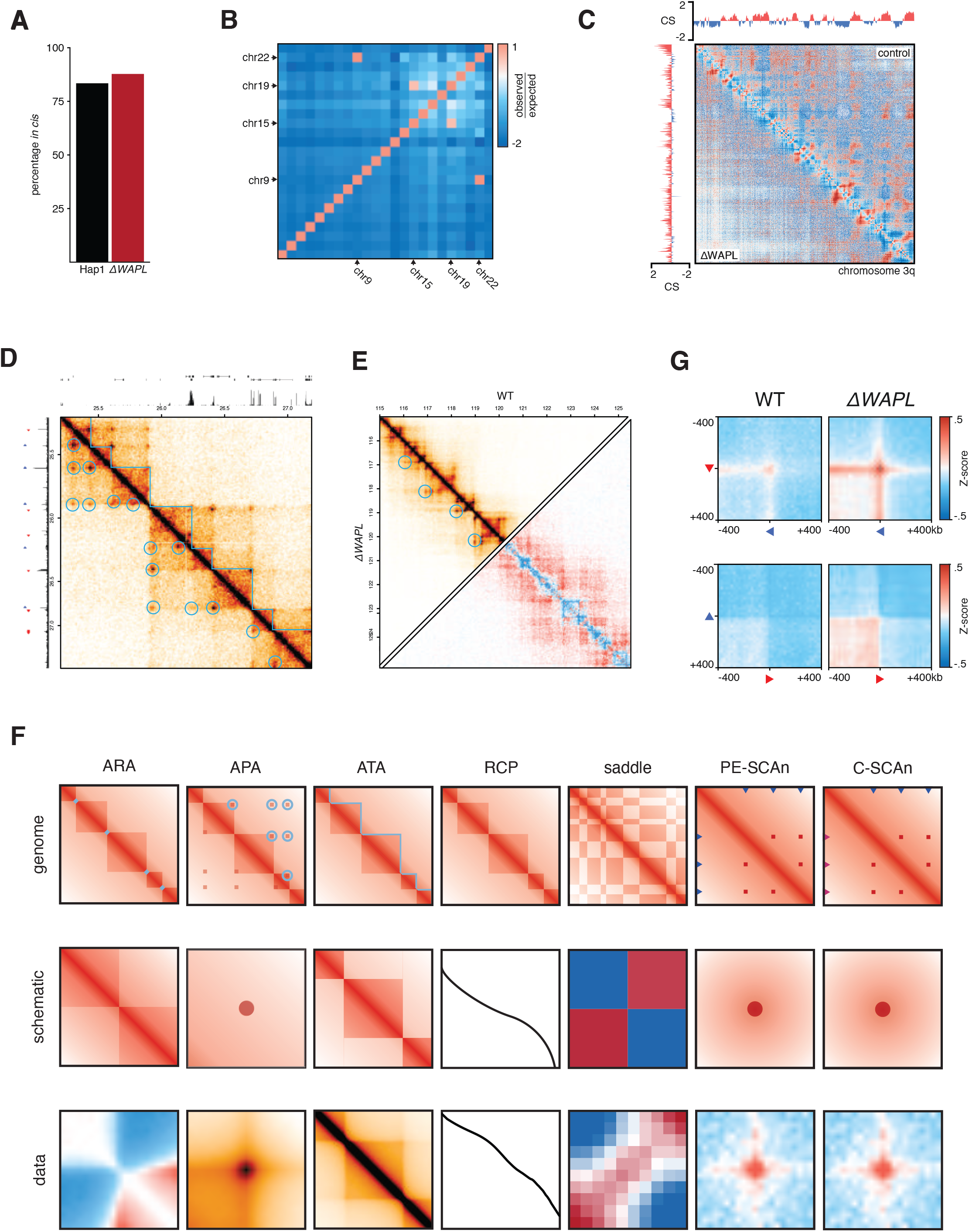
The GENOVA-package contains a complete suite of tools for Hi-C analyses. A) Quantification of the percentage contacts in *cis* of WT and *ΔWAPL* made with the `cis_trans` tool. B) Enrichments of contacts between all pairs of chromosomes with `chromosome_matrix`. Both the reciprocal 9-22 translocation and the addition of a fragment of chromosome 15 on chromosome 19 lead to a high enrichment-score. C) Whole-arm chromosome matrices with compartment-scores for WT (top right) and *ΔWAPL* (bottom left). The matrix can either be the Pearsson-matrix (shown) or the contact-intensity. D) The `hic.matrixplot` tool allows for the plotting of a regions of interest, including annotations. Signal-tracks, gene-models and ChIP-seq peaks can be used for the annotation-tracks above and to the left, while loops and TADs can be plotted on top of the matrix. All annotations can be customised on placement and colour. E) Additionally, a second contacts-object can be added to the bottom-left half of the matrix (top triangle) or can be subtracted from the first contacts-object to produce a differential matrix (bottom triangle). F) Features of the Hi-C data (top) can be summarised with the aggregation-tools of GENOVA (middle) to produce genome-wide averages of the features (bottom). G) C-SCAn of pairwise combinations of CTCF ChIP-seq peaks on forward and reverse binding motifs in convergent (top row) and divergent (bottom row) in WT and *ΔWAPL*.

In studies with translocation-prone (cancer-)genomes, the Hi-C data of sites surrounding the breakpoints will be misleading. The same is true when the reads are aligned to draft genomes that may still contain assembly errors, which can be the case for uncommon model system or strains. In the case of structural variation, the regions around a breakpoint will have increased amounts of —seemingly— *trans*-contacts, which are in reality *cis*-contacts of two translocated pieces of chromosome. In the case of a misassembly, actual wild-type *cis*-interaction will appear as translocations. The result in both cases is the appearance of merged and/or deleted TADs and unexpected changes in compartment-scores. It is therefore recommended that translocated chromosomes are omitted from further analyses. GENOVA can compute the enrichment of *cis*-interactions between chromosomes with `chromosome_matrix`. Moreover, this tool generates an overview-plot for checking for translocations (fig. 2B).

### Tracks and matrices

Hi-C data analysis often focusses around comparing features like TADs and compartments. Identifying the locations of these features first requires that the two-dimensional Hi-C data is reduced to a quantitative linear track. GENOVA provides tools to distil Hi-C into linear tracks on compartment- and domain-level. Aside from calling features on these tracks, users can also use them for matrix-annotation, alignments on regions (e.g. tornado-plots) and viewing in genome-browsers.

To generate a matrix overview for an entire chromosome or chromosome arm (i.e. far-*cis* interactions) we devised the `cis.compartment.plot` function. The resulting plot shows a heatmap of one or two contacts-objects. In the case of two experiments either experiment occupies a triangle in the matrix (top or bottom). The plot can show both the absolute Hi-C signal or the observed over expected (i.e., the distance-dependent average) scores. Above and to the side of the heatmap the compartment-scores are plotted (fig. 2C). This matrix is thus a useful way to get an overview of the far-*cis* landscape and even directly compare two samples

In order to determine A- and B-compartments, users can also generate compartment-scores using a separate function (`compartment_score`). The compartment score is determined by first computing an observed over expected matrix for a chromosome (arm). From this matrix one is subtracted and an eigen decomposition is performed. The first eigenvector of the matrix is multiplied by the square root of the corresponding eigenvalue (31). To ensure that positive values are corresponding to euchromatin, we advise correlating the arm-wise compartment-score to the ChIP-seq data of an active histone mark (e.g. H3K4me1). This can be done from within GENOVA: when this correlation is negative, the compartment-score is multiplied by −1 (36).

Compartments are subdivided in sub-megabase TADs. Two common TAD-level metrics are the directionality index and the insulation score (3, 37). GENOVA includes tools for computing these two separate metrics for TAD-level tracks. It goes beyond the scope of this study to discuss the various downsides and benefits of either method, for a more detailed discussion we refer the reader to (38). These tracks can be used to call TADs and align on genomic features, like genes or precomputed TAD-boundaries (supp. fig. 1A).

The insulation score reflects the differences of contact density of every Hi-C bin with its surrounding bins (37). Briefly, the `insulation_score` tool uses a sliding window to compute the average signal intensity per Hi-C bin. This score is then divided by the genome-wide average signal to produce the insulation-score. To plot the Hi-C matrix and the corresponding insulation score, users can call `plot_insulation`. At the boundary between two TADs there is a clear dip in the insulation score. This feature is exploited in the `call_TAD_insulation` tool to call TAD-boundaries at local minima.

The second TAD-level track, the directionality index, quantifies the bias between upstream and downstream interactions for each Hi-C bin. This score is low just upstream of a TAD-boundary and high just downstream of a TAD-boundary, as has been extensively described by Dixon et al. (2012). The `direct_index` tool will, in short, average the signal in a set region upstream and downstream of a Hi-C bin. Afterwards it is normalized in a similar matter as computing the χ^2^ metric, where a score of zero means that there is no bias. A bin where this score suddenly crosses zero means that interactions are biased up- or downstream, which is the case at TAD-boundaries.

Plotting Hi-C data in user-specified regions in combination with genomic features or data can be done with `hic.matrixplot` (fig. 2D). It accepts multiple sources of annotations: linear features such as ChIP-seq peaks and gene information can be plotted above and to the left of the matrix. TADs and chromatin loops are are plotted over the Hi-C matrix heatmap. Furthermore, linear tracks in bigwig- and bedgraph-format can be plotted to add quantitative information about protein-DNA interactions or gene expression. Two samples can be plotted in a mirrored fashion alongside the diagonal (i,e, the top and bottom triangles of the matrix) or the difference can be plotted by subtracting one experiment from the other (fig. 2E).

### Chromosome-level analyses

The relative contact probability can be used to investigate distance-dependent contact frequencies (1, 39). Because chromosomes are subject to polymer physics (31) the probability of two regions on a chromosome interacting in 3D decreases as function of the linear distance. When comparing two Hi-C experiments, a change in the relative contact probability (RCP) in the 1-5Mb range is indicative of a change in contacts in TAD-level, for example. Moreover, Gassler et al. (40) have shown that the derivative of the RCP can be used to estimate the average extruded loop size. The RCP tool in GENOVA can be used to calculate genome-wide RCP score or for a user-defined set of regions or chromosomes. In addition to the standard methods of plotting the RCP decay as a function of distance for every sample, GENOVA offers the option to compute the fold-change over a control sample (15) (supp. fig. 1B).

While the RCP can give insight into the far-*cis* interactions, it is not designed to reveal changes in the strength of the compartmentalisation, which is measured as the degree in which A and B compartments segregate in the nucleus. For this, users can use the de saddle-tool, which is based on the work of Imakaev et al. (31). In brief, the tool first stratifies each genomic bin on the quantiles of the compartment score. The number of quantile bins can be chosen by the user. Pairwise interactions are then allocated to the combination of their compartment-score quantiles. Next, it computes the average of the observed over expected Hi-C score for each quantile-combination. This results in a N_quantile_xN_quantile_ sized matrix, which can be visualised as a heatmap, a so-called saddle plot. The name of this method comes from the fact that the resulting plot resembles a saddle, with strong interactions at A-A and B-B and weaker interactions between A and B.

A related measure is the compartment strength, which computes the strength of compartmentalisation as the product of the observed over expected (O/E) scores in A/A and B/B (i.e. within compartment) interaction bins divided by the square of the O/E scores in the A/B (i.e. between compartment) interaction bins. A score of one means that the within-compartment interactions are as common as between-compartment, whereas a higher score means that within-compartment interaction are more prevalent.

### Data aggregation

*De novo* TAD and loop calling relies on a sufficiently sequenced dataset (at least 10^8^ reads for the human genome). However, when data is sparse (e.g. less than 25 million reads) we can still extract meaningful information from these datasets through the aggregation over genomic features. GENOVA can perform several forms of aggregation analysis. (fig. 2F).

GENOVA has a family of tools for aggregating contacts at features of interest, like peaks, loops and TADs. Users can aggregate the regions around one-dimensional features (e.g. ChIP-seq peaks or transcriptional start sites, TSS) at the diagonal with the Aggregate Region Analysis (ARA). Since subtle changes can be obscured by the high contact-intensity of the diagonal, the tool computes an observed over expected score. This expected score is generated by calculating the same aggregate matrix for the same features, but shifted 1Mb downstream, and averaging per distance. The Aggregate Peak Analysis (`APA`) averages the signal surrounding the pixels making up the loop taking by default a region 21 bins around the feature (fig. 2F). To aggregate TADs, the `ATA`-tool extracts both the regions of interest (i.e., TADs), including the regions up- and downstream of half of the TAD-size. We average the matrices, after resizing through bilinear interpolation of the individual matrices, to show the average contact-distribution of all TADs and their surroundings (fig. 2F).

All three aggregation-tools have customisable thresholds for the sizes of the feature and its surrounding region to include. Setting the feature-size threshold allows for stratification of specific sizes, such as large versus small loops, but can also be used to remove features that are not in the expected size-range. Users can set a threshold on pixels (i.e., interaction-bins) with extreme values, which are often considered outliers. When a pixel has a higher signal than the threshold, the pixel-intensity will be set to the value of the threshold. This approach keeps all features, regardless of outliers, but limits the influence of the outliers on the final average. Afterwards, the visualise- and quantify-methods allow for comparisons between feature-sets and samples.

Another possibility to visualise aggregates is to generate a tornado-plot, in which the enrichment is plotted for every individual feature (i.e., loop). We calculate the enrichment of each feature with the pixels surrounding it with the same distance (supp. fig. 1C). Afterwards, we sort and k-means cluster the features—both the samples to sort on and the number of clusters can be set. As is the case for all discovery-objects and plots in GENOVA, the output of the tornado contains the raw data, which allows users to further analyse these features, stratified on the clustering.

Aside from Hi-C features, GENOVA also enables the aggregation of contacts between two one-dimensional regions, like ChIP-seq peaks (fig. 2F). PE-SCAn (24) creates virtual loop anchors by combining pairs of features within certain distance-thresholds and calculates the enrichment. C-SCAn is an extension of PE-SCAn and allows multiple sets of peaks (e.g. enhancers and promoters or positively and negatively oriented CTCF motifs). It then creates virtual loops based on combinations of these sets. The discovery-object of PE-SCAn and C-SCAn can be visualised and quantified in the same way as the APA, ARA and ATA.

### Genome editing and cell culture

Hap1 cells were cultured in Iscove’s Modified Dulbecco’s Medium (IMDM) supplemented with 10% FCS (Clontech), 1% Penicillin/Streptomycin (Invitrogen) and 0.5% UltraGlutamin (Lonza). Hap1 SA1 and SA2 knock-out cells were generated using gRNA’s targeting SA1 exon 2 (ACTACTGCCCATTCCGATGC) and SA2 exon 3 (TGATGACCATTCATTCGGTT), which were cloned into PX330. Cells were transfected with PX330 and pDonorTia containing a puromycin resistance gene. Clones were selected using puromycin (2 μg/μl). Colonies were screened for the loss of SA1 and SA2 using PCR and western blot analysis. Used antibodies for the western blots were ab4457 (SA1) from Abcam, 158a (SA2) from Bethyl, sc365189 (WAPL) and sc13119 (HSP90) from Santa Cruz. Rad21 immunofluorescence was performed with Millipore 05-908 (Rad21) antibodies in 1:250 dilution.

### Hi-C from Hap1 SA1 and SA2 knockouts

We performed in-situ Hi-C, as described in Haarhuis et al. (2017). Sequencing was done on the HiSeq X sequencing platform and mapped with hic-pro 2.11.1. We performed loop calling with HiCCUPS 1.9.9.

Previously published Hap1 data (WT and *ΔWAPL*) was included in this manuscript (7). We used both the ice-normalised Hi-C matrices and generated z-normalised matrices during loading. TAD- and loop-calls from the same manuscript were also included. To compare our results to a different cell line, we downloaded the sequencing-reads and juicer-files for the siControl, siSA1 and siSA2 of MCF10A from Kojic et al. (2018). We mapped the reads with hic-pro 2.11.1 (33) to the hg19 reference genome with default settings.

## Results

### Performance and benchmarking

We have developed GENOVA on the premise that it combines all the key Hi-C analysis tools for the most common Hi-C data formats. To illustrate that contacts-objects from different formats can be compared in GENOVA, we mapped the data of Kojic et al. (2018) with HiC-pro and compared it to .hic files mapped with TADbit and converted with Juicer-tools. The relative contact probabilities between the two formats are similar for both siSA1 and siSA2 (supp. fig. 1D). This shows that the different formats give nearly identical output and that these different outputs can be compared inside GENOVA.

Because Hi-C maps are often large and complex datasets, the speed of these tools is key to many of the analyses. Therefore, we use key-based binary searches(41), which has the benefit that the speed of the analyses is no longer linearly proportional to the number of regions queried (41). To test the performance of our method, we performed an Aggegrate Peak Analysis on Hap1 Hi-C data of Haarhuis et al. (2017) with both GENOVA and Juicer(23). Our analysis showed that, irrespective of resolution, the absolute increase in calculation time is less with more loops queried in our implementation (supp. fig. 1E). These results indicate that GENOVA’s implementation of region-lookups is robust and quick enough to handle large queries.

Aggregation enables the gathering of information from dataset that have a higher level of sparsity. To investigate how sparse the data can be and still be used in aggregation-analyses, we downsampled the HAP1 data of Haarhuis et al. (2017). The RCP analysis shows that there is little to no deviation of the full dataset up to 90Mb at 1 million reads (supp. fig. 1B). Both the APA and ATA show good signal-to-noise, even at 5 million reads—twenty percent of the output of a current Illumina MiniSeq (supp. fig. 1F,G). These results indicate that aggerate analyses can be faithfully performed on low-coverage datasets.

### C-SCAn and loop clustering

In GENOVA we have implemented two novel tools, C-SCAn and loop clustering. The first is an extension of the previously published Paired-End Spatial Chromatin Analysis (PE-SCAn) method(42), that aggregates of all pairwise combinations of a genomic feature such as gene promoters or super enhancers(43). C-SCAn builds on this by performing aggregation of pairwise combinations of two different genomic features, for instance gene promoters and distal enhancers, but excluding the homotypic pairwise combinations. We tested our method by aggregating over combinations of forward and reverse oriented CTCF binding sites. Our analysis showed, as expected, that there was a clear increased contact frequency between CTCF binding sites in a convergent orientation (fig. 2G). This contact frequency was further increased in the absence of WAPL, consistent with the observation that cohesin is bound more stably to DNA(7). Note that the C-SCAn function allows the user to analyse genomic features in a specific direction, like with the forward and reverse CTCF sites, or in a direction agnostic manner, as with promoters and enhancers. The C-SCAn function is a powerful new method to elucidate features that shape the 3D genome.

A powerful method to visualise ChIPseq data is a heatmap of the signal around, for instance, peaks, also referred to as tornado plots. We realised that, for obvious reasons, no such method existed for Hi-C data. We have therefore developed a method that selects diagonals from the Hi-C matrix that overlap with specific points in said matrix, such as chromatin loops or putative chromatin loops, represented as a one-dimensional array of values. These arrays can be stacked together in a heatmap, similar to ChIPseq tracks. Visualization of the heatmap enables the assessment of global versus specific changes in loop changes (supp. fig. 1C). The organisation of the loop data into a matrix also enables the user to perform *k*-means clustering, to identify specific subsets of loops (discussed in more detail below). These are two additions to a roster of analysis tools that can be used to analyse Hi-C. Below we will use these tools to analyse the role of different cohesin variant in nuclear organisation.

### Differing far-cis landscapes of cohesin^SA1^ and cohesin^SA2^

The cohesin-complex has been shown to play a major role in the formation of CTCF-anchored loops and contacts within TADs (6). There are two variants of the complex, containing either the SA1 (STAG1) or SA2 (STAG2) homologs, that are suggested to have specialised functions (fig. 3A) (15, 16, 30). To elucidate the differences of cohesin^SA1^ and cohesin^SA2^ with regard to genome organisation, we made knock-outs of either SA1 or SA2 by inserting a puromycin resistance cassette in-frame in HAP1 cells (supp. fig. 2A). We confirmed the knockouts by PCR (supp. fig. 2B) and western blot (fig. 3B). We refer to these knock-out lines as ΔSA1 and ΔSA2.

**Figure 3:**
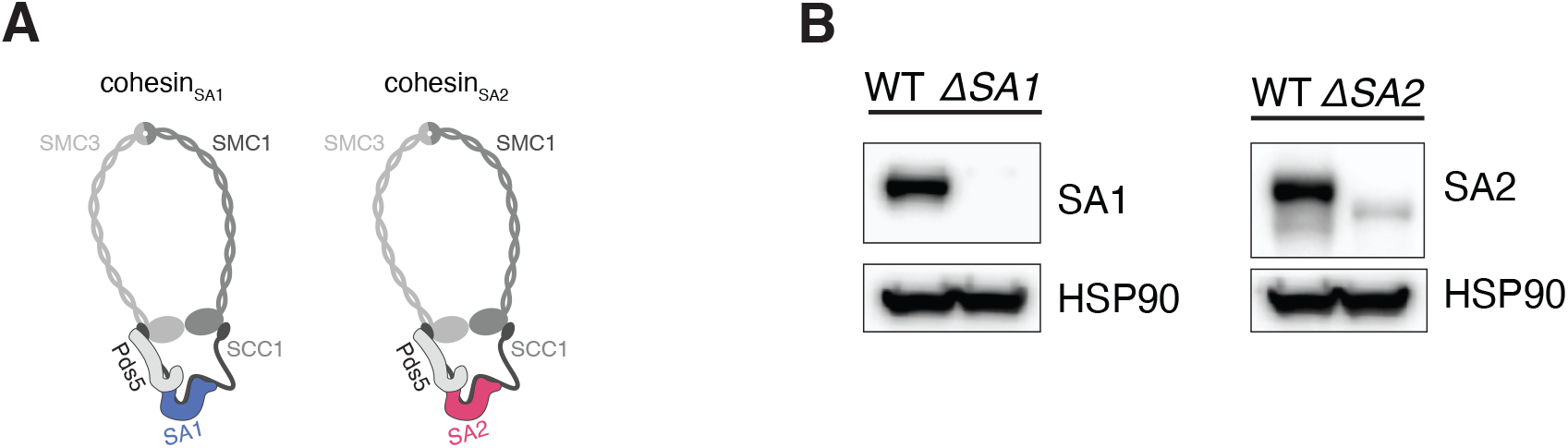
Generation of Hap1 SA1 and SA2 knockouts. A) The two cohesin-variants differ in their SA subunits. B) Western blot analysis confirms SA1 knockout in ΔSA1 cells and ΔSA2 knockout in ΔSA2 cells.

To reveal the effects of knocking out SA1 or SA2 on chromosome organization, we generated high-resolution Hi-C maps. When inspecting whole chromosome-arms, we saw that the two knockouts had different effects on the intrachromosomal interaction landscape. In ΔSA1 cells there were more far-*cis* interactions, indicated by the stronger “plaid”-pattern in the Hi-C map. On the other hand, in ΔSA2 cells there are more interactions at the sub 5Mb-scale, which can be seen as a stronger diagonal (fig. 4A, supp. fig. 3A). This difference was confirmed in the relative contact probability (RCP) plots, where the ΔSA2 has increased interactions in the close-*cis* range (1-10Mb), compared to the WT. The ΔSA1 cells show a general increase in contacts compared to WT for regions more than 5Mb apart. (fig. 4B, supp. fig. 3B). We found that the technical replicates show extremely similar distributions, and thus combined the replicates in all subsequent analyses (supp. fig. 3B). Our results indicate that cohesin^SA1^ and cohesin^SA2^ affect chromosome organization differently.

**Figure 4:**
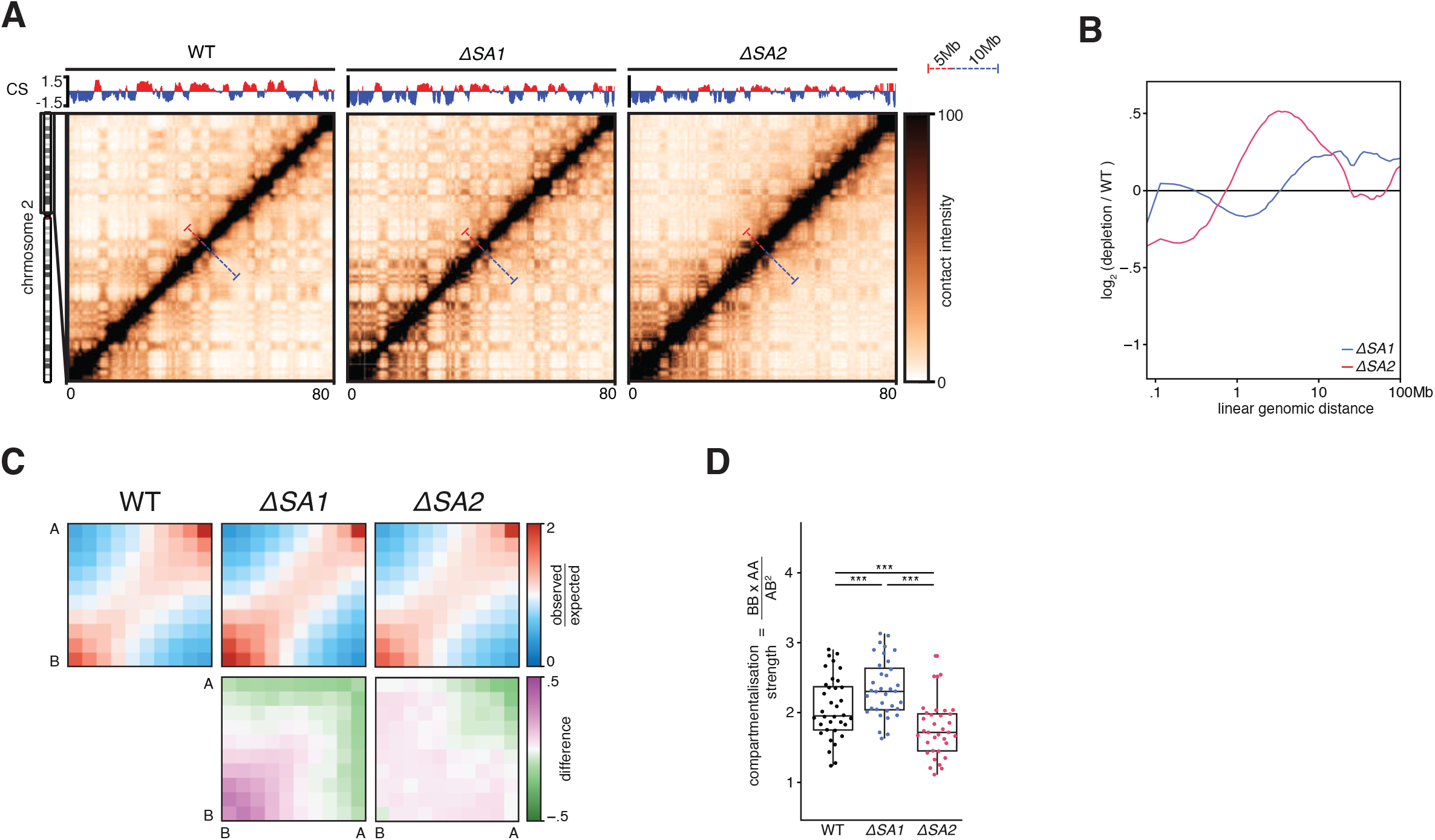
Far-*cis* differences between the cohesin-variants. A) Hi-C matrices of chromosome 2p of wild-type, ΔSA1 and ΔSA2. Compartment-scores are plotted on top. Bars in matrices denote 5mb and 10mb distances in red and blue, respectively. B) Relative contact probabilities compared to wild-type in log2-space, with blue denoting ΔSA1 and red denoting ΔSA2. C) Saddle-plots (top) and differential saddles (bottom), with purple denoting more interactions in the sample compared to the wild-type. D) Boxplot of the compartmentalisation-strength per chromosome-arm (dots). *** indicates paired t-test p < .005.

The observation that ΔSA1 has increased far-*cis* interactions compared to ΔSA2 brings up an interesting possibility that cohesin^SA1^ inhibits compartmentalisation (i.e. more intra-compartment contacts) to a larger extent than cohesin^SA2^. This difference in compartmentalisation can already be seen in the compartment-score tracks of figure 4A: the amplitude of the B-compartment score (blue) is increased in the ΔSA1 compared to both the WT and ΔSA2. Since a higher compartment-score amplitude is an indication of more homotypic compartment interactions (i.e. between two A compartment bins or two B-compartment bins), we quantified these differences genome-wide. To this end, we generated saddle-plots to quantify the amount of self-interaction of A- and B-compartments (31, 44). These plots show that ΔSA1 has increased B-B (and less A-B) interactions compared to control (fig. 4C). This can be further quantified using the compartment strength(31), which corresponds to the proportion of intra- versus inter-compartment contacts and is calculated for every chromosome arm separately (31). We found that the ΔSA1 overall has significantly stronger compartmentalisation, while ΔSA2 has weaker compartmentalisation, compared to wild-type (fig. 4D). These results show that cohesin^SA1^ and cohesin^SA2^ differ in their propensity to restrict compartmentalisation.

### Cohesin^SA2^ promotes intra-TAD contacts

Depletion of the cohesin loading/extrusion factor Scc2/Nipbl or loss of the cohesin loading factor SCC4/MAU2 leads to an increase in compartmentalisation, whereas cohesin stabilization on DNA reduces compartmentalization (7, 45). From this it has been postulated that cohesin loops actively counter compartmentalisation (46). We thus investigated chromosome organisation at the level of chromatin loops. TADs are thought to be an average representation of cohesin-mediated chromatin loops. Therefore a difference in loop formation activity should be visible at this level of resolution. We indeed observed a striking difference in TADs between both cohesin-variants (fig. 5A, supp. fig. 4A). In ΔSA2, TADs show an increased signal at the edges (i.e. corner peaks) and diminished intra-TAD signal. We used the TAD-calling tool in GENOVA, which is based on the insulation score(37), to identify TAD-boundaries in all samples. The number of TAD-boundaries between WT and ΔSA1 was similar, whereas the ΔSA2 has a decreased number of boundaries (fig. 5B). Furthermore, the overlap of TAD-boundaries between ΔSA1 and WT is three-fold higher than ΔSA2 versus WT. These results suggest that cohesin^SA2^ plays a role in the formation of intra-TAD contacts, which in turn leads to a stronger insulation of TADs.

**Figure 5:**
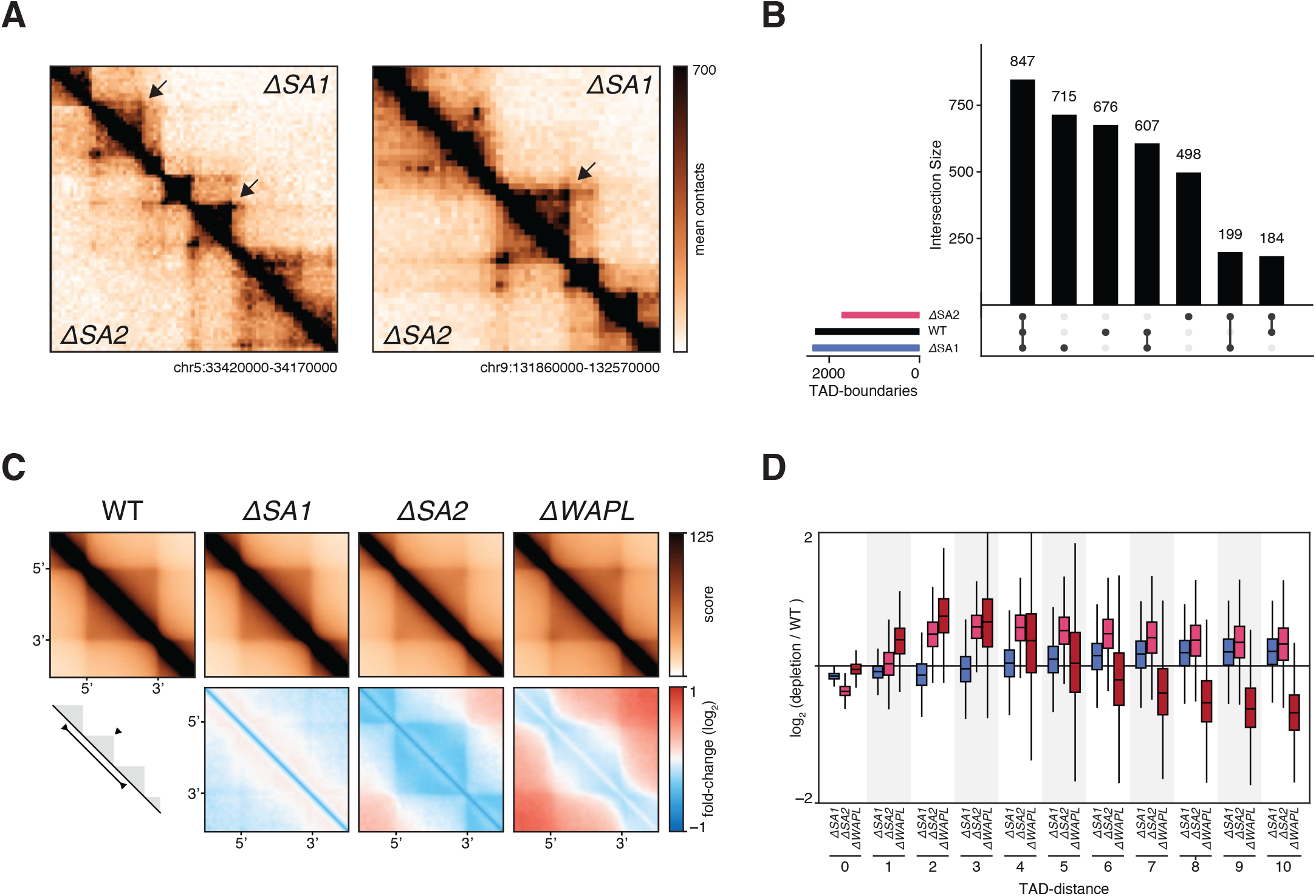
Cohesin^SA1^-only cells have diminished intra-TAD contacts. A) Snapshots of two regions on chromosome 5 and chromosome 9, showing ΔSA1 in top-right and ΔSA2 in bottom-left triangle. B) Intersections of called TAD-boundaries in wild-type, ΔSA1 and ΔSA2. C) Aggregate TAD analysis of Hap1 TADs in wild-type, ΔSA1, ΔSA2 and ΔWAPL (top). Differential ATA compared to wild-type (bottom), which blue denoting loss of interactions in the specific sample. D) TAD-neighbour analysis: interactions between TADs, stratified on the number of TADs in between, compared to wild-type.

Our observations regarding TADs in ΔSA2 cells were reminiscent of loop formation in ΔWAPL. Stabilisation of cohesin by loss of WAPL also leads to more-pronounced corner peaks at TAD boundaries, and fewer intra-TAD interactions. To further explore the consequences on TADs, we performed an Aggregate TAD Analysis (ATA) on TADs called in Haarhuis et al. (2017). The ATA shows that the aforementioned loss of intra-TAD contacts in ΔSA2 is found genome-wide (fig. 5C, supp. fig. 4B). Moreover, the quantification of the ATA indicates that ΔSA1 have increased intra-TAD off-diagonal contacts (supp. fig. 4C). Loss of SA2 by RNAi in MCF10A cells (13) results in a similar phenotype (supp. fig. 4D).

The similarity at the level of TADs between ΔSA2 and ΔWAPL prompted us to investigate the contacts over boundaries. The `intra_inter_TAD` tool in GENOVA enables this comparison a systematic manner. As shown previously(7), there are more interactions between (maximal 5) neighbouring TADs in the ΔWAPL, while the intra-TAD score is decreased (fig. 5D). On the other hand, intra-TAD contacts are decreased even more in ΔSA2 cells and inter-TAD score increases as far away as 10 TADs. These findings again suggest that cohesin^SA2^ is required for intra-TAD contacts.

### Cohesin^SA1^ creates longer CTCF-anchored loops

FRAP experiments have recently shown that cohesin^SA1^ is more stably associated with chromosomes than cohesin^SA2^ (14). We hypothesize that a longer residence time of cohesin on chromatin leads to the formation of longer loops. One way to measure this is to investigate a feature of Hi-C maps called “stripes”, which are formed at CTCF sites and thought to be a manifestation of one-sided loop extrusion by cohesin. We measured stripe formation in our Hi-C data by performing an ARA on CTCF-sites with a specific orientation (supp. fig. 5A). We observed a pattern that is reminiscent of insulation consistent with the function of CTCF. Furthermore, a clear stripe pattern is found in the direction of the CTCF site. In ΔSA1 cells the stripe signal decays more rapidly compared to the wild-type (supp. fig. 5B). In contrast, the ΔSA2 cells show hardly any decay compared to the wild-type over the distances we measured. In addition to this, we also see an increase in contacts upstream of the CTCF-site in cells that only have cohesin^SA1^ (supp. fig. 5B). This increase of upstream contacts at CTCF-sites is in line with the presence of bidirectional anchors due to loop-extension, as anchors of extended loops are combinations of CTCF-loops themselves (47).

Upon further inspection of the Hi-C matrices we indeed observed loops over larger distances in the ΔSA2 cells, which only have cohesin^SA1^ (fig. 6A). To systematically investigate these differences, we called loops with HICCUPS and calculated the size-distribution per genotype (fig. 6B). We find that the average loop-size is increased in the ΔSA2 from 410kb to 500kb. Conversely, in the ΔSA1 the average loop length is decreased to 320kb. The ΔSA2 specific longer loops connect loop anchors already found in wild-type (fig. 6A). We systematically analysed this using a function in GENOVA that enables the calculation of average contact frequency between extended loops, that are formed between the 5’ and 3’ anchors of loops called in wild-type cells. The APA for these extended loops showed that ΔSA2 cells show an increase in the contact frequency (fig. 6C), which is reminiscent of results we previously observed for ΔWAPL cells (7). We also observed this in the data of Kojic et al. (2018), where the SA2-depletion showed an increased signal at extended loops (supp. fig. 5C). To exclude that the effect on loop length that we are seeing is an indirect effect of lower WAPL levels, we performed Western blot analysis. This confirmed that the WAPL protein level was unaffected (supp. fig. 5D). Together, these analyses support the notion that the stability of a cohesin-variant on chromatin determines the length of the loops that can be produced.

**Figure 6:**
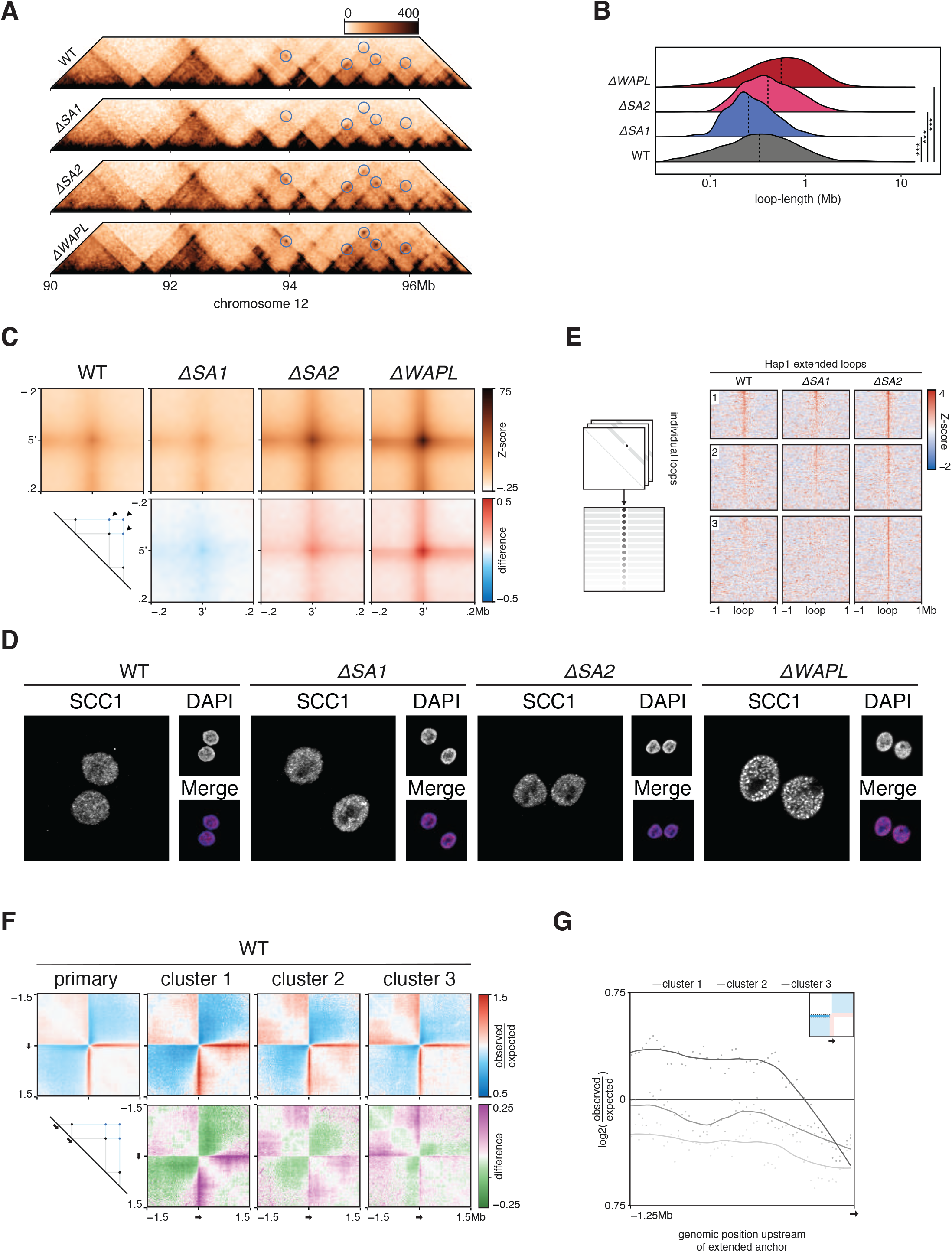
Extended loops in ΔSA2 are formed at bidirectional loop-anchors. A) Pyramid-plots of wildtype, ΔSA1, ΔSA2 and ΔWAPL at chromosome 12. Predicted loop-extensions, based on wild-type anchors, are indicated in blue circles. B) Length-quantification of loops called by HICCUPS in wild-type, ΔSA1, ΔSA2 and ΔWAPL. Dashed line denotes median. C) Aggregate peak analysis of the predicted extended loops in wild-type and the three knockouts (top). Differential plots comparing knockouts to wild-type are shown in the bottom row, where red indicates an enrichment in the knockout. D) Immunofluorescence of DNA-bound SCC1, showing the vermicelli-phenotype in ΔWAPL. E) The aggregate tornado-plot extracts the signal around and at every individual loop visualises them as a heatmap, with a loop at every row (left). A *K=3* clustered tornado on the APA-discovery object of figure 6C. Cluster 3 harbours ΔSA2-specific extended loops. F) Aggregate region analysis on wild-type data, using upstream anchors of all loops (primary) and those of the extended loops from the clusters found in figure 6E. G) Quantification of the upstream regions from the ARA of figure 6F (left) and of the complimentary analysis of the downstream anchors (right).

Loss of WAPL also leads to increased stability of cohesin on DNA and an increase in loop length. This is accompanied by a striking ‘vermicelli’ chromosome phenotype in which a thread-like staining of cohesin is seen. Because of the increased loop size in ΔSA2 we investigated whether the vermicelli phenotype is also found in our ΔSA2 cells. To this end, we stained the cohesin subunit SCC1 in WT, ΔSA1, ΔSA2 and ΔWAPL cells. Whereas the ΔWAPL cells showed a clear vermicelli phenotype, the ΔSA2 cells lack vermicelli chromosomes (fig. 6D). These results show that, although the absence of WAPL and SA2 correlate with an increase in loop size and the formation of extended loops, further differences in cohesin stability likely determine whether vermicelli chromosomes are formed (see Discussion).

### Extended loops form at bidirectional anchors

Because both ΔSA2 and ΔWAPL cells show extension of loops, but result in different chromosome organization at the ultrastructural level, we looked in more detail at the extended loops in these different genotypes. To quantify and cluster the underlying loops of the APA, we used the aggregate tornado tool. Running this tool on our data showed that there are three clusters, of which cluster 3 (containing 674 pairwise sites) has a strong enrichment in the ΔSA2 only (fig. 6E). This enrichment shows that cohesin^SA1^ can form extended loops when cohesin^SA2^ is absent at previously identified loop-anchors.

Casual observation of extended loops in figure 6A already revealed that not all loop anchors have the same propensity to form extended loops. To determine whether there are any predictive features for extension in the ΔSA2, we compared the signal in the WT-cells of these anchors in the different clusters, as well as the complete set of WT-anchors. We performed an ARA on the 5’ anchors in the wild-type Hi-C data (fig. 6F). The anchors of all three clusters show the expected stripe in the downstream direction (i.e. the direction of the called loop). Surprisingly, however, we observed a difference in contact enrichment in the upstream direction. The quantification of the signal upstream of the anchor showed that cluster 3 anchors have a stronger upstream signal and showed stripe-like behaviour in the opposite orientation (fig. 6G, left). When performing the same analyses for the downstream anchors, we also see that cluster 3 anchors have the strongest signal outside of the loop (fig. 6G, right). These results suggest that bidirectional anchors (which have both up- and downstream loops in the wild-type) are more likely to gain extended loops in the ΔSA2.

## Discussion

Here we present GENOVA, an R package that combines the most important Hi-C data analyses and which can be run on commodity hardware. GENOVA has powerful visualization tools for a suite of analyses, ranging from relative contact probability plots to compartmentalization analyses and aggregations of TADs and loops. While visualization is an important aim in Hi-C data analysis, GENOVA also provides tools to quantify the underlying data for specific analyses. For instance, when the user runs an analysis to check the average contact frequency for a set of loops, the result can be visualized. However, the relevant pixel information can also be extracted using quantification tools. These data can then be visualised and analysed with one of the many visualisation and statistical tools available in R. Specifically for this reason the package does not contain options to automate null-hypothesis testing. Due to that the sheer number of possible tests and comparisons we leave it up to user to choose the statistical test that matches their data type. We are confident that running the `quantify`-tool on the discovery-objects of the aggregations, provides the user with enough options to pursue these tests outside of GENOVA.

The aggregation analyses also enable the analyses of more sparsely sequenced datasets. The costs of sequencing Hi-C matrices to kilobase resolution can be quite daunting, especially when replicates are involved. By performing aggregation analyses, relevant information can be extracted from datasets that are sequenced at relatively low depth. Importantly, this also opens the door for performing analyses on replicate experiments, which are now often combined into a single dataset to boost the visualization. Obviously, these analyses work only for perturbations that have a general effect on 3D genome organization. For perturbations that affect only a handful of loops in the genome, deeper sequencing will still be required.

A number of tools have been developed that enable the browsing of Hi-C data such as Juicebox(23) and HiGlass(48). These tools also enable adding one-dimensional tracks for ChIPseq and RNAseq data, for instance. Although GENOVA does not allow interactive browsing of Hi-C data, it does offer the creation of publication-ready Hi-C matrix plots that can be annotated with genomic features and genomic data tracks. A powerful suite of tools that has an overlapping feature set with GENOVA is HiCexplorer(49). This is a command line tool that is written in Python, we command structure that is similar to the popular deeptools package(50).There is a large number of dependencies, which makes this package difficult to install on an operating system such as Windows. Because GENOVA is written in R it is largely platform agnostic and we have confirmed installation on Linux, Windows and MacOS. With the increasing popularity of R with in the genomics and broader life sciences community we believe that GENOVA can serve as an important go-to package for Hi-C data analysis for experimentalists and bioinformatics-specialists alike.

### Cohesin variants differently contribute to 3D genome organisation

Here we studied the roles of variant cohesin-complexes on chromosome organisation. We made knockouts in Hap1 cells of either SA1 or SA2, generated Hi-C data and performed analyses in GENOVA. We find that cohesin^SA1^ produces longer loops, while cohesin^SA2^ is biased towards shorter loops. The stronger compartmentalisation in cohesin^SA2^-only cells is consistent with a decrease in loop extrusion, as suggested by (7, 11, 46).

The differences in loop length are consistent with recent FRAP experiments that surveyed the residence time of the two cohesin variants by measuring cohesin association with DNA in the absence of either SA1 or SA2(14). Cohesin^SA1^ was shown to have a longer chromatin residency time, which was suggested to result in longer extrusion and longer loops. Interestingly, co-depletion of CTCF with SA2 diminished cohesin^SA1^ residence time to wild-type levels, indicating that cohesin binding to chromatin is stabilised by CTCF. If CTCF leads to long-term stabilisation of cohesin the observed differences in loop length may also be the result of differences in extrusion kinetics between the cohesin variants. If cohesin^SA2^ would be slower to extrude, fewer cohesin complexes would reach a distal CTCF site and ultimately result in cohesin complexes stably associated with DNA. Recent advances in *in vitro* single molecule imaging experiments of cohesin-mediated DNA extrusion (51, 52) offer an exciting opportunity to measure these parameters. Alternatively, measuring loop formation kinetics using Hi-C following mitosis(53) or rapid reconstitution of RAD21 proteins levels(6) in an SA1 or SA2 null background should be able to address this question.

Finally, it has been speculated (based primarily on the loss of intra-TAD contacts) that cohesin^SA2^ plays a role in enhancer-promoter interactions, while cohesin^SA1^ is thought to be responsible for looping together CTCF binding sites (13, 15, 16). Our current results suggest that this distinction is too strict, as we show that CTCF-anchored loops are still present in the ΔSA1 cells This is further supported by the fact that other reports also show that CTCF-loops are still present in SA1-depletion lines (13–15, 30). It should be noted that cohesin’s CTCF binding pocket is conserved in both SA1 and SA2 (54). It therefore is likely that CTCF can bind and regulate both cohesin variants. Our current results show that the different cohesin variants contributed differently to genome organization. Varying the levels of SA1 and SA2 relative to each other could therefore be an important mechanism to regulate genome organization and gene expression. How these variants contribute to or counteract the function of the other variant in the wild-type situation will be an important question for the future.

### Vermicelli versus extrusion

As described previously and again in this study, SCC1-staining during WAPL depletion leads to a thread-like distribution of cohesin in interphase nuclei as measured by immunofluorescence, known as the vermicelli phenotype (58). This —and the fact that loops become extended— had been attributed to the increased stability of cohesin onto chromatin (7). In the ΔSA2 cells we found extended loops, but not a vermicelli phenotype. An explanation could be the model above, in which the cohesin^SA1^-only cells have increased cohesin-stability, but not enough compared to ΔWAPL to form sufficient numbers of loop-collisions to be visible as vermicelli. Multi-contact analyses are necessary to determine whether in the absence of SA2 loop collisions are indeed not formed (12). Further research into the formation of loop-extension and the vermicelli phenotype is also needed to provide evidence for this model or uncoupling of the two phenotypes.

Concluding, we propose a model in which cohesin-variants have differing loop formation kinetics, which leads to the changes in nuclear architecture that we observe. This points towards another layer of chromatin-regulation: balancing of the loops formed between specific anchors to ensure a proper chromatin landscape.

## Availability

GENOVA is an open source software package in the GitHub repository http://www.github.com/dewitlab/GENOVA.

## Accession numbers

Data has been deposited at GEO under accession GSE160490.

## Acknowledgement

We thank the members of the division of Gene Regulation for helpful discussions and support, Marijne Schijns, Lucas Kaaij and Ángela Sedeño Cacciatore for testing and improving GENOVA, the NKI Genomics Core Facility for sequencing, and the NKI microscopy facility for help with imaging.

---

**Supplementary Figure 1:**
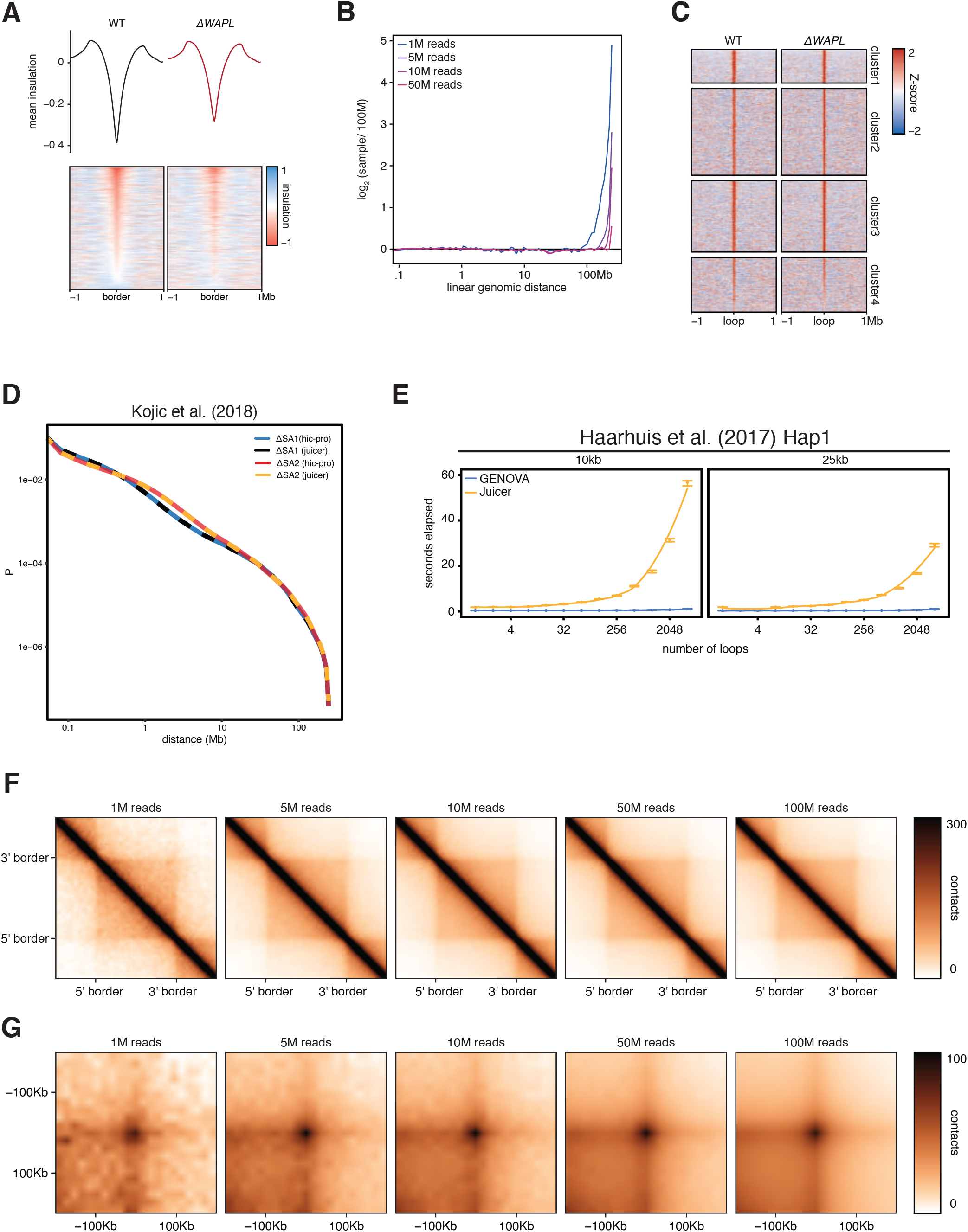
GENOVA can produce tornado-plots of tracks and aggregations and produces robust results. A) Calculating and aligning the insulation score of multiple samples can be done with the `insulation_score` and `insulation_tornado` tools. B) Log-fold changes in RCP between the downsampled Hi-C matrices and the full high-depth dataset. C) Tornado-plots of an APA-discovery object, containing primary loops of Haarhuis et al. (2017), with K=4. D) RCP-outputs of siSA1 and siSA2 from Kojic et al. (2018), from either the hic-pro or juicer input. E) Number of loops surveyed in an APA versus the time taken for both 10kb and 25kb. Blue line denotes APA tool from GENOVA; yellow line denotes APA from juicer_tools.jar. F,G) ATA and APA of WT Hap1, subsampled to various levels of sparsity.

**Supplementary Figure 2:**
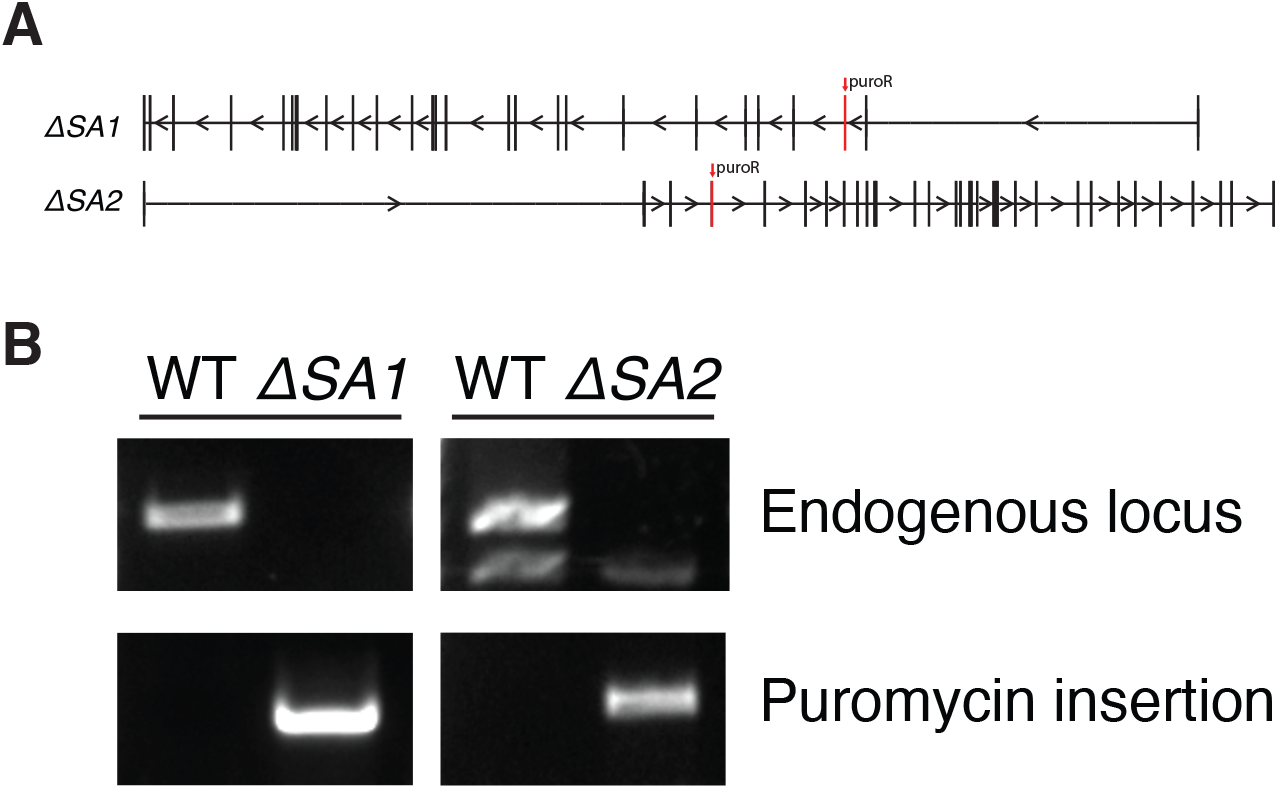
Knockout-strategy of Hap1 ΔSA1 and ΔSA2. A) A puromycin-cassette is inserted by CRISP-cas9 in the third and fourth exon of SA1 and SA2, respectively. B) PCR confirms the loss of the endogenous locus and the gain of the puromycin-cassette at the targeted sites.

**Supplementary Figure 3:**
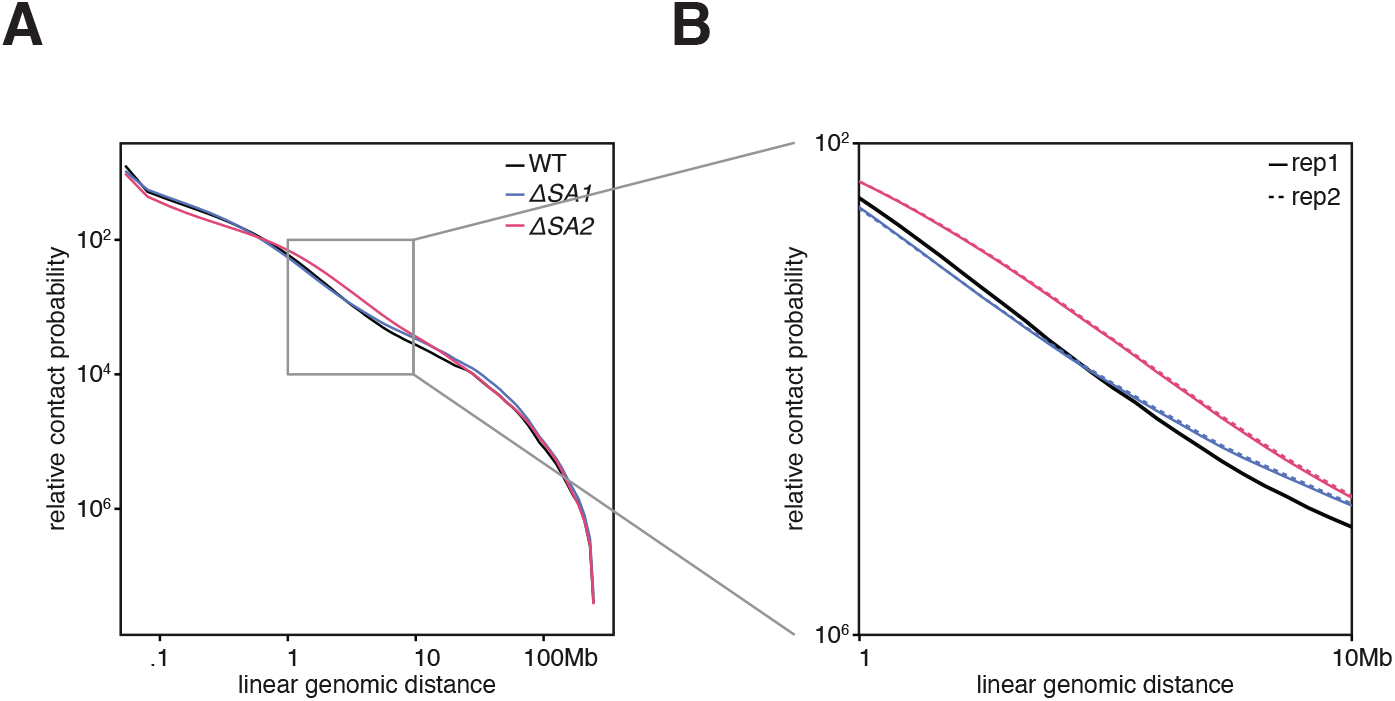
Contact-distributions of ΔSA1- and ΔSA2-replicates show strong similarity. A) Relative contact probabilities of wildtype (black), ΔSA1 (blue) and ΔSA2 (red). Square denotes 1-10mb distance-range. B) Zoom of 1-10Mb distances, showing the individual replicates. Replicate 1 in solid line and replicate 2 in dashed line.

**Supplementary Figure 4:**
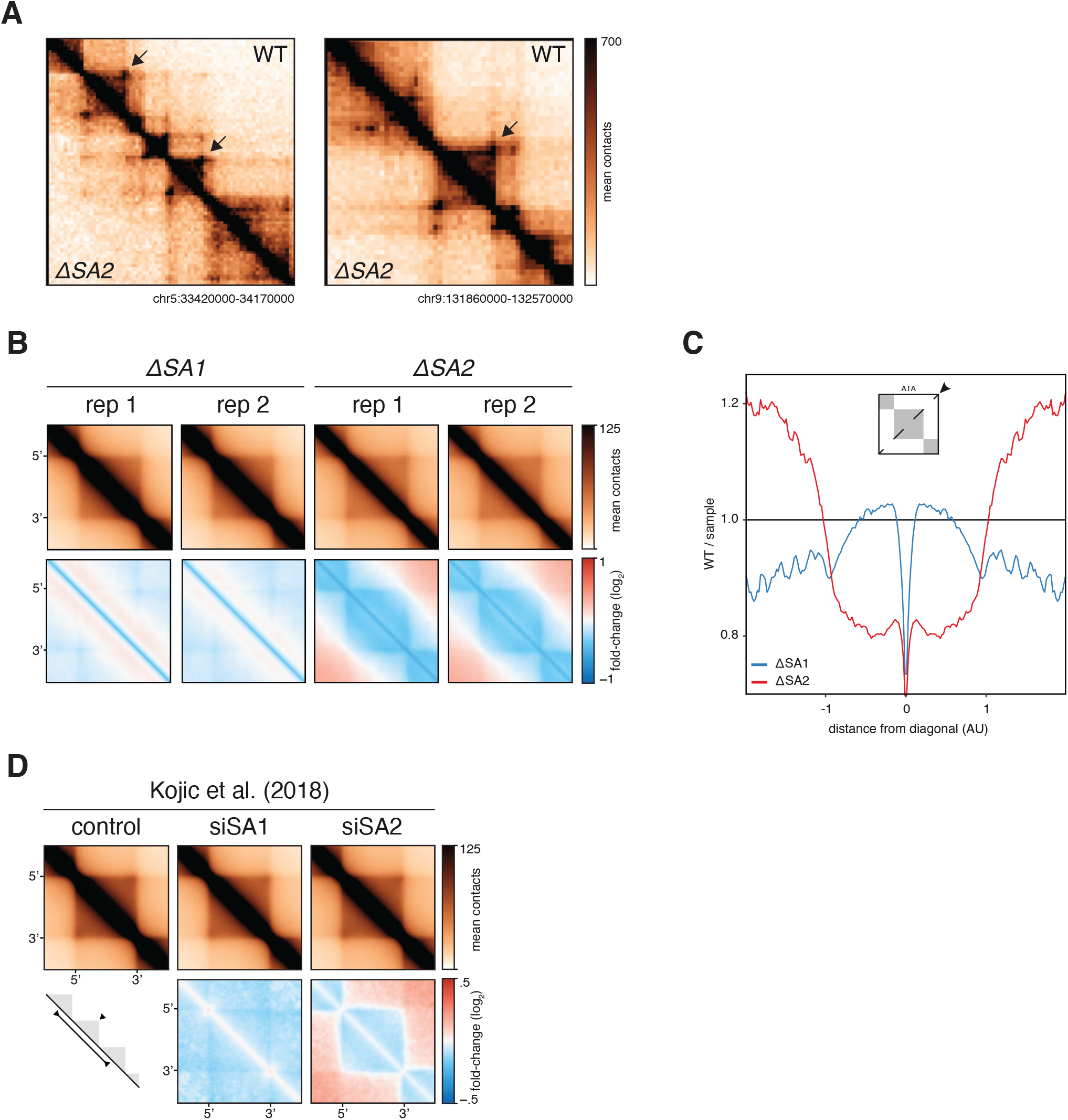
The loss of intra-TAD interactions is reproducible in SA2-knockouts and - depletions. A) Snapshots of two regions on chromosome 5 and chromosome 9, showing wild-type in top-right and ΔSA2 in bottom-left triangle. B) Aggregate TAD analysis of Hap1 TADs in the separate replicates of ΔSA1 and ΔSA2 (top) and compared to wild-type (bottom). C) Quantification of the second diagonal in the ATA in B. D) Aggregate TAD analysis of MCF10A TADs in control, siSA1 and siSA2 data of Kojic et al. (2018).

**Supplementary Figure 5:**
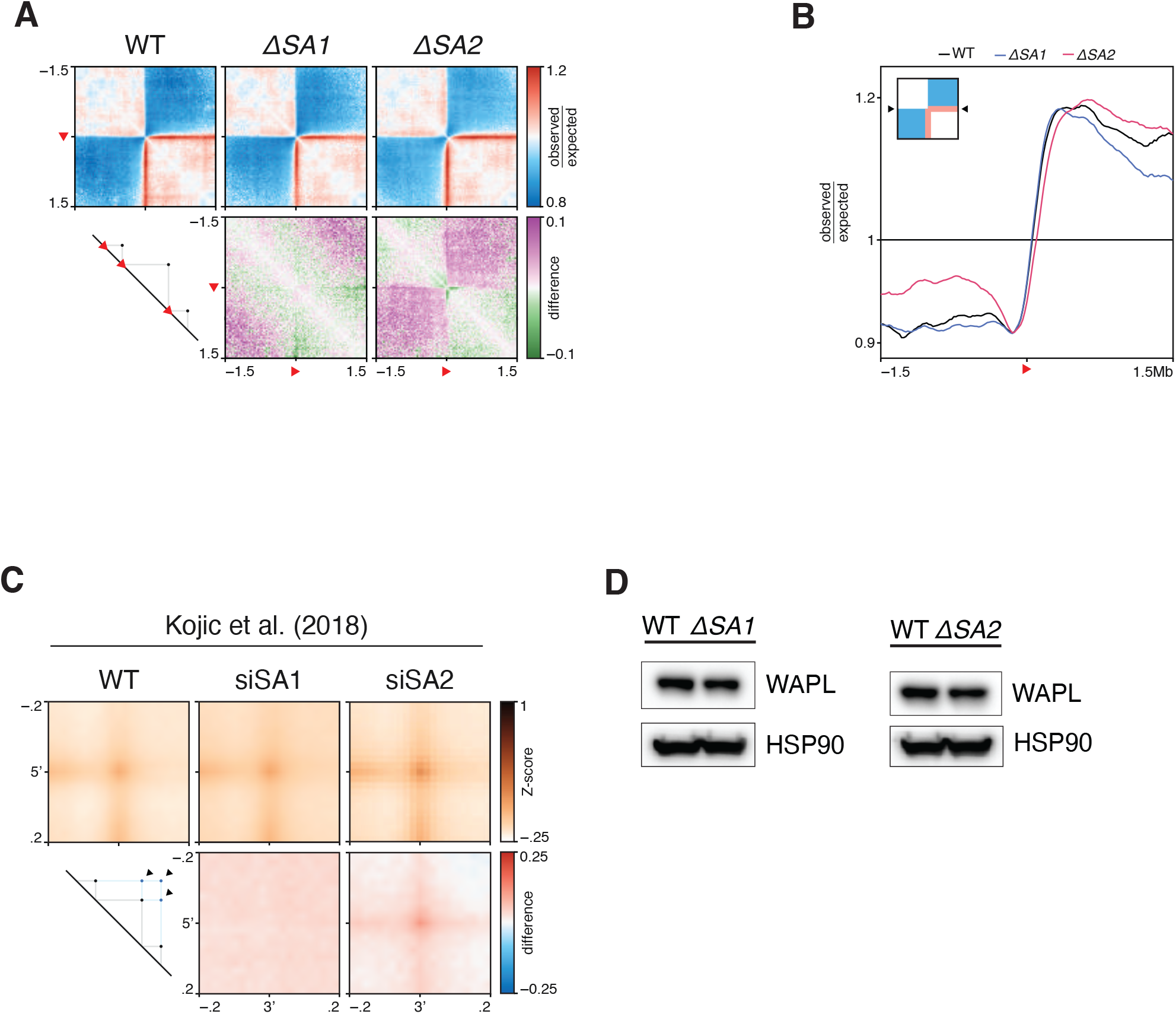
CTCF-stripes are increased in ΔSA2. A) Aggregate region analysis of wild-type, ΔSA1 and ΔSA2 on CTCF-motifs in the forward orientation. B) Quantification of the ARA on forward CTCF sites of supplementary figure 5A. C) APA of predicted extended loops in the data of Kojic et al. (2018). D) Western blot analysis confirms that WAPL levels are unaffected in ΔSA1 and ΔSA2.

